# Characterization of *cis*-prenyltransferase complexes in guayule (*Parthenium argentatum*), an alternative natural rubber-producing plant

**DOI:** 10.1101/384149

**Authors:** Adam M. Lakusta, Moonhyuk Kwon, Eun-Joo Kwon, Solomon Stonebloom, Henrik V. Scheller, Dae-Kyun Ro

## Abstract

Guayule (*Parthenium argentatum*) is a perennial shrub in the Asteraceae family and synthesizes a high quality, hypoallergenic *cis*-1,4-polyisoprene (or natural rubber; NR). Despite its potential to be an alternative NR supplier, the enzymes for *cis*-polyisoprene biosynthesis have not been comprehensively studied in guayule. Recently, implications of the protein complex involving *cis*-prenyltransferases (CPTs) and CPT-binding proteins (CBPs) in NR biosynthesis were shown in lettuce and dandelion, but such protein complexes have yet to be examined in guayule. Here we identified four guayule genes – three *PaCPTs* (*PaCPT1-3*) and one *PaCBP*, whose protein products form PaCPT/PaCBP complexes. Co-expression of both *PaCBP* and each of the *PaCPTs* could complemented the dolichol (a short *cis*-polyisoprene)-deficient yeast, whereas the individual expressions could not. Microsomes from the *PaCPT/PaCBP-expressing* yeast efficiently incorporated ^14^C-isopentenyl diphosphate into dehydrodolichyl diphosphates. Furthermore, co-immunoprecipitation and split-ubiquitin yeast 2-hybrid assays using PaCPTs and PaCBP confirmed the formation of protein complexes. Of the three *PaCPTs*, transcriptomics analysis indicated that the protein complex formed by PaCPT3 and PaCBP is likely to be the key component in guayule NR biosynthesis. The comprehensive analyses of these PaCPTs and PaCBP here provide the foundational knowledge to generate a high NR-yielding guayule.

## Introduction

Natural rubber (NR) is an isoprene biopolymer, chemically defined as *cis*-1,4-polyisoprene, and it is used to manufacture a number of medical and industrial products (Cornish, 2001; van Beilen and Poirier, 2007). NR is best known for its prized physical properties, such as elasticity, abrasion resistance, heat dispersion, and hysteresis, which have special importance for the products used in medical and heavy-duty industries. Despite significant advances in polymer science, no material can so far fully replicate the qualities of NR (Mooibroek and Cornish, 2000). The key feature that distinguishes the NR from other synthetic polymers is that NR is a stereo-regular polymer. The isoprene monomeric units of NR are condensed entirely in *cis*-1,4-configuration in average Mw of ~1 million Da. Such uniform stereo-structures are theorized to permit NR to form an extremely lengthy spring-like biomolecule, whereas synthetic rubbers are not exclusively stereo-regular (Morton, 2013). These properties, therefore, make NR a strategically important, irreplaceable commodity in modern manufacturing industries.

NR is synthesized in >7,000 plant species (Rivano *et al*., 2013). Nonetheless, the rubber tree (Para rubber tree or the Brazilian rubber tree; *Hevea brasiliensis*) from the Euphorbiaceae family is an almost exclusive NR-producing plant species. The rubber tree synthesizes NR on rubber particles in latex, the cytoplasmic content of the specialized laticifer cells (Cornish *et al*., 1999). The supply of NR solely depended on the rubber trees, mostly (>90%) cultivated in Southeast Asian countries (van Beilen and Poirier, 2007). During World War II, the supply route of the NR from Southeast Asia was blocked by the Japanese Empire, and in search of alternative NR-producing plants, American and Russian researchers identified guayule (*Parthenium argentatum*) and the Russian dandelion (*Taraxacum kok-saghyz*), respectively, as these plants can also synthesize high quality NR (Mooibroek and Cornish, 2000; van Beilen and Poirier, 2007). However, the research activities to develop alternative rubber plants ceased at the end of World War II.

Guayule is a perennial desert shrub native to the northern states of Mexico, but it also grows in the southern states of Texas, New Mexico, and Arizona in the USA. Guayule is unique in its ability to grow in some of North America’s most arid lands, receiving as little as 25-38 cm of rainfall annually (Rasutis *et al*., 2015). In addition to having little need of water, guayule is known to have dry weights ranging from 4–12% in NR (Cantú *et al*., 1997). Unlike rubber tree, guayule does not have laticifers, but instead, synthesizes NR in the parenchyma cells in stem (Kajiura *et al*., 2018).

The NR biosynthetic mechanism has not been fully elucidated yet, and researchers have focused on the *cis*-prenyltransferase (CPT) class of enzymes, as these enzymes are able to catalyze the condensation of isopentenyl diphosphate (IPP) onto a priming molecule in an exclusively *cis*-configuration (Teng and Liang, 2012). The prokaryotic CPT family enzyme, undecaprenyl diphosphate synthase (UPPS), catalyzes the condensations of IPP onto farnesyl diphosphate (FPP) to form primarily C55 undecaprenyl diphosphate (eight IPP molecules condensed to one FPP) (Shimizu *et al*., 1998). Its monophosphate form, undecaprenyl phosphate, is a vital lipid molecule that mediates sugar transfer for peptidoglycan cell-wall biosynthesis in prokaryotes. The crystal structure study demonstrated UPPS to be a soluble, homo-dimeric enzyme localized in the prokaryotic cytosol (Fujihashi *et al*., 2001). Similarly, a homologous eukaryotic CPT enzyme, dehydrodolichyl diphosphate synthase (DDS), catalyzes the analogous *cis*-condensation reaction to form C80 - C120 *cis*-polyprenyl diphosphate, which is converted to dolichyl phosphate by a phosphatase, polyprenol reductase, and dolichol kinase (Denecke and Kranz, 2009). The resulting dolichyl phosphate serves as a sugar carrier molecule for the post-translational modification of proteins in all eukaryotes. Therefore, both UPPS and DDS are essential metabolic enzymes, and lesions on these genes are lethal (Sato *et al*., 1999; Zhang *et al*., 2008). Interestingly, the primary and tertiary structures of CPTs display no similarities to those of trans-prenyltransferases, indicating that **cis*-* and trans-prenyltransferases do not share a common origin and have independently evolved for distinct stereo-selective catalytic reactions (Fujihashi *et al*., 2001; Lu *et al*., 2009).

Although comprehensive biochemical data for the prokaryotic CPTs have been documented in the literature, *in vitro* biochemical characterizations of eukaryotic CPTs has been relatively scarce in the literature until recently. This discrepancy was first resolved by the discovery of the human CPT binding partner, NogoB receptor. Human CPT forms a hetero-protein complex with the NogoB receptor and becomes active in dolichol biosynthetic pathway (Harrison *et al*., 2011). Similar complexes comprised of CPT and its binding protein homolog were also found in lettuce, tomato, and Arabidopsis for dehydrdolichyl diphosphate biosynthesis (Brasher *et al*., 2015; Qu *et al*., 2015; Kwon *et al*., 2016). The CPT binding proteins, including the NogoB receptor, were named differently in each species. In this work, they are referred to as CBP (CPT-binding protein). CBP is only found in the eukaryotic lineage from yeast to humans, but interestingly, not in prokaryotes (Brasher *et al*., 2015; Qu *et al*., 2015). CBP displays low sequence homology to CPTs, forms a complex with CPT on the endoplasmic reticulum (ER), and lacks all conserved motifs required for *cis*-prenyltransferase activity. Therefore, it is believed that CBP interacts with, activates, and tethers CPT onto the eukaryotic ER, but does not function directly in polymerization.

Intriguingly, RNA interference studies showed that analogous CPT-CBP complexes are also involved in NR biosynthesis in lettuce and dandelion (Epping *et al*., 2015; Qu *et al*., 2015). However, *in vitro* reconstitution solely using microsomal CPT and CBP recombinant proteins failed to produce *cis*-polyisoprenes longer than dolichol (Qu *et al*., 2015). It is worth noting that lettuce encodes two pairs of *CPT-CBP* genes - one for the dehydrodolichyl diphosphate biosynthesis within primary metabolism, and the other for NR biosynthesis for the specialized (or secondary) metabolism in latex (Qu *et al*., 2015). Therefore, at least in lettuce and related species, such as dandelion, it is evident that the NR biosynthesis has functionally diverged from the primary dolichol metabolic pathway.

While the evidence to support the model for CPT-CBP complex formation in the NR biosynthesis exists in lettuce and dandelion, both species are closely related to each other. Each of these species belongs to the same Cichorieae tribe of the Cichorioideae subfamily within the Asteraceae family. Outside lettuce and dandelion (Cichorieae tribe), the rubber tree and guayule are the only plants that are commercially cultivated for NR production. Of these two, guayule (*P. argentatum*) is from the same Asteraceae family as lettuce and dandelion, but it comes from a different subfamily and tribe (Asteroideae and Heliantheae, respectively), whereas the rubber tree (*H. brasiliensis*) belongs to an entirely different family, Euphorbiaceae. Recently, studies of CPT and CBP from the rubber tree suggested that its CPTs interact with its CBP (Yamashita *et al*., 2016).

The Heliantheae (for guayule) and the Cichorieae (for lettuce and dandelion) tribes are distantly related within the Asteraceae family as they share a common ancestor from ~39 million years ago (www.timetree.org). Considering such a divergence between guayule and lettuce/dandelion, along with guayule’s commercial potential to produce NR in arid regions, we conjecture that in-depth molecular studies of guayule *CPTs* and *CBP* can provide the knowledge necessary to help develop guayule as an alternative NR-producer. Guayule is transformable (Ponciano *et al*., 2018), and utilization of its biomass via bio-refinery process has been studied (Orts *etal*., 2016). Isolated rubber particles from guayule have been used for biochemical studies (Cornish and Siler, 1995), but guayule *CPT* and *CBP* have not been investigated at the molecular level.

In this work, *CPT* and *CBP* homologs were identified from guayule, followed by the investigation of their biochemical activities, interactions, and expression patterns. In addition, the findings from the guayule CPT and CBP studies are discussed in the context of the current knowledge of NR biosynthesis.

## MATERIALS and METHODS

### Plant materials and growth conditions

Guayule plant seeds (PI478663) were obtained from the U.S. Department of Agriculture (USDA). Seeds were soaked in fresh water for 4 days before germination, and potted to a mixture of 35% autoclaved sands and 65% potting soil. The germinated plants were grown in a growth chamber under conditions of 16 h light, 8 h dark, at 25°C.

### RNA isolation, gene isolation, and phylogenetic analysis

Two months old guayule plants were ground in liquid nitrogen using mortar and pestle, and 100 mg of powder was mixed with 1 mL of Trizol (Invitrogen), and total RNA was extracted according to the manufacturer’s protocol. First strand synthesis of cDNA was performed using 1 μg of the extracted RNA (leaf and stem) as template, combined with the M-MuLV reverse transcriptase (NEB) and anchored oligo dT_22_ (IDT). The synthesis of cDNA was performed according to NEB’s provided protocol. *PaCPT1* was amplified from guayule cDNA by primers 1/2 and cloned into the *pGEMT-easy* vector (Promega) using TA/blunt ligase mix (NEB). *PcCPT2/3* and *PaCBP* were synthesized after yeast codon optimization. Phylogenetic and molecular evolutionary analyses were conducted using *MEGA* ver. 6 (Tamura *et al*., 2013). All sequences were retrieved from NCBI database, and rubber tree *CBP* was identified from published sequencing data (Tang *et al*., 2016).

### Yeast complementation assay and *in vitro* biochemical assay

Details of yeast complementation, *in vitro* enzyme assay, and reverse-phase thin layer chromatography were previously described (Kwon *et al*., 2016). *PaCPT1-3* were amplified using primers 3/4, 5/6 or 7/8, respectively, and the amplicons were cloned into p414-GPD using SpeI/SalI, SpeI/XhoI or BamHI/ClaI sites, respectively. *PaCBP* was cloned into p415-GPD using SpeI/XhoI sites. The cloned vectors were introduced to a yeast strain deficient in both copies of CPTs, *rer2Δ∷HIS3 srt1Δ∷KanMX*[pRS316-RER2], described in Kwon *et al.* (2016). This strain is referred to as *rer2*Δ *srt1*Δ hereafter, and Ro lab yeast stock number is ROY13.

### Split ubiquitin membrane yeast two-hybrid (MYTH) and ß-galactosidase assay

MYTH assays were performed according to the published methods (Gisler *et al*., 2008; Snider *et al*., 2010). The bait plasmid (*PaCBP-Cub-LexA-VP16*) was generated by ligating the *PaCBP* amplified with primers 9/10 into pAMBVα using StuI/NcoI sites. For the prey plasmids, *PaCPT1-3* were amplified with primers 11/12, 13/14 or 15/16, and cloned into pPR3C-NubG in SpeI/BamHI (*PaCPT1*) or BamHI/ClaI (*PaCPT2 & 3*) to yield *PaCPT-HA-NubG* constructs. *PaCPT1-3* were also amplified with primers 17/18, 19/20 or 21/22 and cloned into pPR3N-NubG in BamHI/SalI (*PaCPT1*) or BamHI/ClaI (*PaCPT2 & 3*) to generate *NubG-HA-PaCPT* constructs. The bait and prey plasmid pair was co-transformed into the Y2H reporter yeast strain, THY.AP4. The control prey vectors pOST1-NubG and pOST1-NubI were also co-transformed with the bait vector. Individual transformed-yeast colonies were cultured in 30°C overnight, and the cells were spotted on SC-Leu-Trp (SC-LT) and on SC-Leu-Trp-Ade-His (SC-LTAH) containing 3-amino-1,2,4-triazole (3-AT) at concentrations of 5, 10, or 25 mM. Spotted plates were incubated for 3 days at 30°C to observe cell growth.

The strength of protein interactions was quantified using β-galactosidase assay with *O*-nitrophenyl-β-D-galactoside (ONPG) as the substrate. From SC-LTAH selection plates, the positive colonies were cultured overnight in SC-LTAH medium without 3-AT. Cells from the overnight culture were inoculated to 4 mL of SC-LTAH medium and cultured for 5 hours. The cells were harvested at 9,000 × g for 3 min at 4 °C and resuspended in 1 mL of Z buffer (60 mM Na_2_HPO_4_, 40 mM NaH_2_PO_4_, 10 mM KCl, 1 mM MgSO_4_ and 50 mM β-mercaptoethanol, pH 7.0), 20 μL of chloroform and 20 μL of 0.1% SDS. After vortexing, the mixture was pre-incubated for 5 min at 28 °C. The reaction was initiated by adding 0.2 mL of ONPG (4 mg mL^-1^ in Z buffer) and terminated by the addition of 1 M of Na_2_CO_3_. The mixture was centrifuged at 20,000 × *g* for 10 min, and the absorbance at 420 nm and 550 nm were measured to determine the Miller units (Miller, 1972).

### Cloning and *In vitro* Translation of *PaCPT1-3* and *PaCBP*

DNA constructs for *in vitro* translation were cloned into pT7CFE1-3xFLAG (constructed using pT7CFE1-NHa) for *PaCPT1-3* and pT7CFE1-NHa (ThermoFisher Scientific) for *PaCBP*.pT7CFE1-3xFLAG was generated by replacing HA tag in pT7CFE1-NHa with 3xFLAG. The 3xFLAG (atggactacaaagaccatgacggtgattataaagatcatgatatcgattacaaggatgacgatgacaag) was amplified using primer sets 23/24, 23/25, and 23/26 sequentially using the IRES region in pT7CFE1-NHa as a template. The final PCR product containing partial IRES sequence and 3xFLAG was digested by KpnI and BamHI and ligated to corresponding restriction sites in pT7CFE1-NHa. Synthetic PaCPT1 and PaCPT2 (GenScript, NJ, USA) were ligated to BamHI and XhoI sites in pT7CFE1-3xFLAG. PaCPT3 was amplified from *P. argentatum* leaf cDNA using a primer set, 27/28. The PaCPT3 PCR product was cloned into pT7CFE1-3xFLAG via BamHI and XhoI restriction sites. Synthetic PaCBP (GenScript, NJ, USA) was amplified using a primer set 29/30 and digested with BamHI and XhoI, followed by ligation to corresponding sites in pT7CFE1-NHa. A control construct, pT7CFE1-NHa-GFP for co-immunoprecipitation described below, was generated by cloning *GFP* sequence into pT7CFE1-NHa. *GFP* was amplified from pCFE-GFP (ThermoFisher Scientific) using a primer set 31/32, containing BamHI and XhoI, respectively, followed by ligation to corresponding sites in pT7CFE1-NHa. The purified plasmid preparations of all IVT constructs were cleaned up further by ethanol precipitation, followed by resuspension in nuclease-free water to a concentration of 500 ng μL^-1^ prior to IVT. IVT was carried out using 1-Step Human Coupled IVT Kit (ThermoFisher Scientific), following the manufacturer’s procedure except for reducing the reaction to half of the recommended amounts. Immediately following incubation of the IVT reaction mixture at 30 °C for 4 hours, 1 μL of the 12.5 μL IVT mixture was saved as input for Western blotting by adding SDS-PAGE loading buffer and storage in -20 °C until use. The remaining IVT mixture was used immediately for immunoprecipitation as described below.

### Co-immunoprecipitation of PaCPT1-3 and PaCBP

For pre-binding of protein G magnetic beads (NEB, ON, Canada) to monoclonal mouse anti-FLAG M2 antibody (Sigma, ON, Canada) or monoclonal rat anti-HA 3F10 antibody (Roche, Quebec, Canada), for each co-immunoprecipitation reaction, 10 μL of the bead suspension was washed (invert 5 times/vortex 5 sec) in 100 μL of room temperature PBS buffer (137 mM NaCl, 2.7 mM KCl, 10 mM Na_2_HPO_4_, 1.8 mM KH_2_PO_4_, pH 7.4) four times and incubated with 0.5 μL of the antibody in 100 μL of cold PBST (PBS, pH 7.4, 0.1% Tween 20) at 4 °C for 16 hours on a Labquake^™^ Tube Rotator. The separation of magnetic beads from solution was achieved by applying magnetic field for 30 seconds on a magnetic stand. To remove unbound antibody, the antibody-immobilized beads were washed (invert 5 times/vortex 5 sec) three times, each in 100 μL of cold IP buffer (150 mM NaCl, 100 mM Tris-HCl pH 7.4, 1 mM EDTA, 10% glycerol, 1% Triton X-100, 0.5% NP-40, 1 mM PMSF). The immobilized beads were resuspended in 500 μL of cold IP buffer, followed by addition of the 11.5 μL of the completed IVT reaction mixture. Co-immunoprecipitation reaction was incubated at 4 °C for 1 hour on the rotator. To remove unbound proteins, the mixture was washed (invert 5 times/vortex 5 sec) five times, each in 700 μL of cold IP buffer. Immunoprecipitated proteins were eluted from the beads by resuspending in 20 μL of 2X SDS-PAGE loading buffer (100 mM Tris-HCl pH 6.8, 4% (w/v) SDS, 20% glycerol, 0.01% (w/v) bromophenol blue, 200 mM β-mercaptoethanol) and incubating at 95 °C for 5 minutes with intermittent gentle flicks. This elution also liberates anti-FLAG and anti-HA antibodies from the magnetic beads as the heavy (50 kDa) and light (25 kDa) chains of the IgG antibodies. After bringing the eluted mixture to room temperature, the mixture was spun down by a brief centrifugation, followed by transfer of the eluate while on a magnetic stand. Half of the eluate was used to run 12% SDS-PAGE followed by Western blotting with mouse anti-FLAG M2 antibody and the other half with rat anti-HA 3F10 antibody. The IVT reaction mixture (1 μL of 12.5 μL total IVT reaction) saved as input was also divided in half to run in SDS-PAGE and Western blotting with anti-FLAG and anti-HA antibodies. SDS-PAGE gel was transferred to PVDF membrane using a Tris-glycine-methanol buffer system, and the antibody hybridization was performed in 5% skim milk TBST (5% skim milk, 50 mM Tris-HCl pH 7.5, 150 mM NaCl, 0.05% Tween 20) at 1:2000 for both anti-FLAG and anti-HA antibodies by overnight incubation at 4 °C. HRP-conjugated anti-mouse and anti-rat secondary antibodies were also hybridized in 5% skim milk TBST at 1:10,000 at room temperature for 1 hour.

### RNA-Seq analysis

For *de novo* transcriptome assembly and differential gene expression analysis, 4 month-old greenhouse-grown guayule plants (AZ-2 cultivar) were transferred to a simulated summer condition in a growth chamber (16 hr daylight, 25°C daytime and 15°C night time temperatures). At 6 months of age, plants were transferred to a simulated winter condition (13 hr daylight, 25°C daytime and 5°C nighttime temperatures) for induction of rubber production. Plants were harvested for analysis at 8 months of age. RNA was extracted from 3 biological replicates each for leaf and stem tissues from induced and control plants using Trizol (Invitrogen) following the manufacturer’s instructions. Following Trizol extraction, RNA samples were treated with DNAase using the Turbo DNA-free kit (Ambion) and further purified using RNeasy column purification (Qiagen). Directional, bar-coded Illumina sequencing libraries were prepared using NEBNext^®^ Ultra^™^ Directional RNA Library Prep Kit (NEB) with the suggestions for size selection of an average insert size of 300-450 bp. Libraries were then pooled and sequenced on Illumina Miseq and HiSeq2500 platforms producing 28 million 300 bp paired end reads and 155 million 150 bp paired end reads, respectively, with an average of 2.4 million MiSeq and 12.9 million HiSeq reads per sample. RNA-seq data was quality-controlled using Trim Galore (Babraham Bioinformatics group) to eliminate adapter sequence contamination and to trim data with a quality score below Q30. Miseq and Hiseq reads were pooled for transcriptome assembly and differential gene expression analysis. The transcriptome was assembled using Trinity assembler version 2.04 set to minimum Kmer coverage of 2 (Haas *et al*., 2013). The initial transcriptome assembly was filtered using a utility included within the Trinity package to eliminate transcript isoforms with low abundance and low coverage. Differential gene expression analysis was performed using EdgeR (Robinson *et al*., 2010). The assembly data are available in the NCBI short read archive under submission number SRP107961.

## RESULTS

### Identification of guayule *PaCPT* and *PaCBP*

Most cultivated guayule (*Parthenium argentatum*) plants for NR production are polyploidy, but the guayule transcriptome assembly from the diploid accession PI478640 is publicly available (Hodgins *et al*., 2014). This data set was retrieved and used in this work as diploid plant makes the analysis of gene family simple. In this transcriptome, 51947 unigenes were assembled from 983076 reads (454 GS FLX Titanium) from four different guayule tissues (root, leaf, flower, and stem). We screened this assembled transcriptome using lettuce *CPT* and *CPT-B/nd/ng Prote/n (CBP)*., formerly referred to as *CPT-L/ke (CPTL)* in our previous study in lettuce (Qu *et al*., 2015). From this analysis, one guayule *CBP* and three *CPT* cDNAs were identified, and these cDNAs were named *PaCBP* and *PaCPT1-3.* Unlike lettuce that has two *CBP* isoforms, one of which displays a laticifer-specific gene expression, guayule does not have laticifer cells and has only a single copy of *CBP.*

### Phylogenetic analysis of *PaCPT1-3* and *PaCBP*

To understand phylogenetic relatedness of *PaCPT1-3* to other known eukaryotic and prokaryotic *CPTs*., a phylogenetic tree was reconstructed using the deduced amino acid sequences from *PaCPT1-3* cDNAs. It has been established that two clades of CPTs (clade I and clade II) are present in plants (Figure 1A). Clade I is comprised of chloroplastic CPTs that likely originated from prokaryotic homo-dimeric CPTs, whereas clade II consists of cytosolic/endoplasmic reticulum (ER) CPTs that interact with CBPs to form hetero–protein complexes, which are then recruited to ER. The CPT clade II is further divided into two sub-clades – one type of CPT for dolichol biosynthesis in the primary metabolism and the other type for NR biosynthesis in the specialized secondary metabolism.

**Fig. 1.**
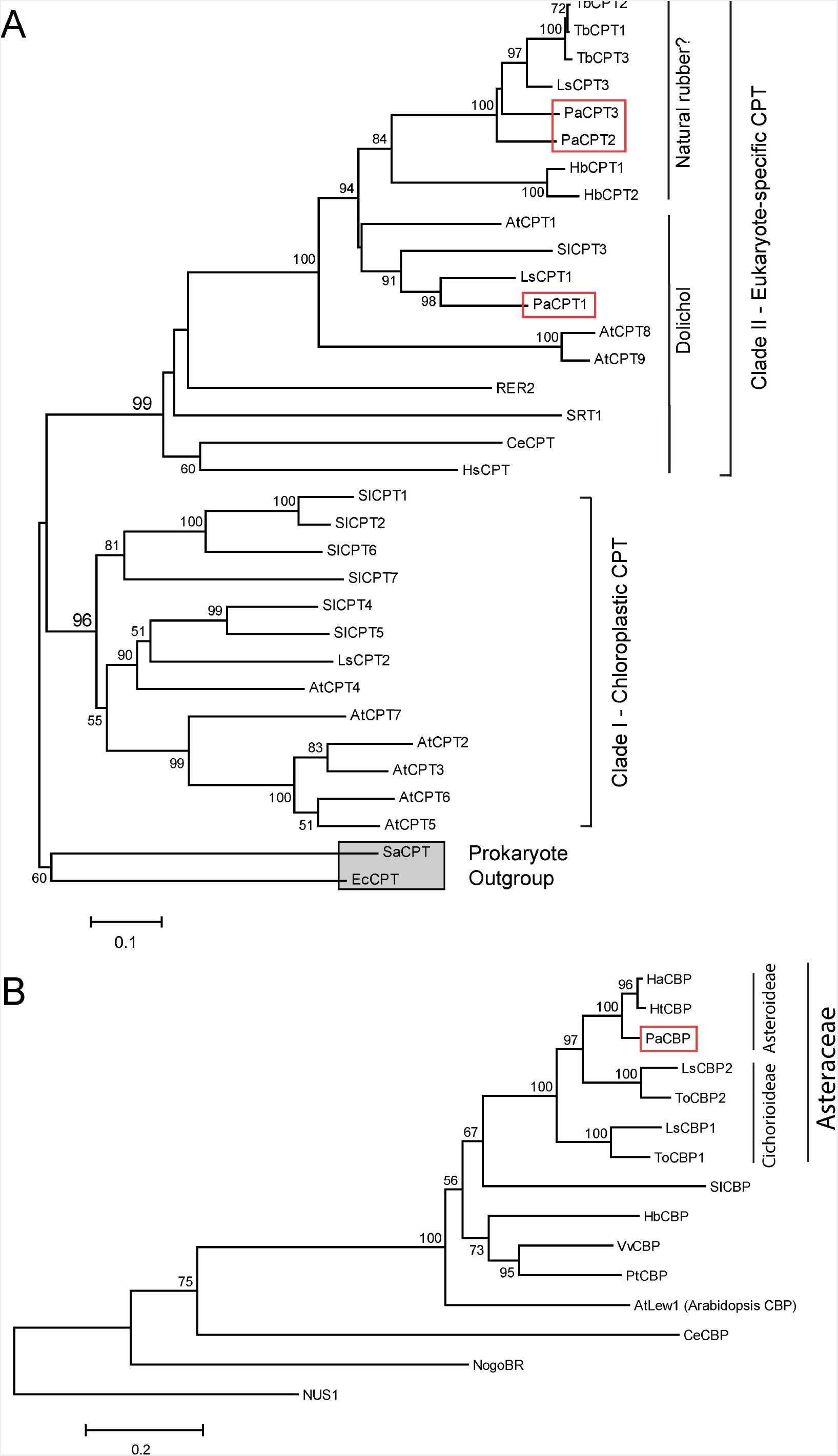
Phylogenetic analysis of CPT and CBP proteins. Phylogenetic trees were created based on the protein sequence similarities between CPTs and CBPs from various prokaryotic and eukaryotic species. Bootstrap values from 1,000 replicates were calculated, and the percentages of replicates are shown in each node only when higher than 50%. Two prokaryotic CPTs (EcCPT and SaCPT) are used as the outgroup and indicated by a grey square. PaCPT1-3 and PaCBP are marked by red boxes. Abbreviations used are: At, *Arab/dops/s thal/ana;* Ce, *Caenorhabd/t/s elegans;* Ec, *Escher/ch/a col/;* Ha, *Hel/anthus annuus;* Hb, *Hevea bras/l/ens/s;* Hs, *Homo sap/ens;* Ht, *Hel/anthus tuberosus;* Ls, *Lactuca sat/va;* Pa, *Parthen/um argentatum;* Pt, *Populus tremulo/des*; Sa, *Staphylococcus aureus;* Sl, *Solanum lycopers/cum;* Tb, *Taraxacum brev/corn/culatum;* To, *Taraxacum off/c/nale;* Vv, *V/t/s v/n/fera; RER2* and *SRT1* are yeast CPTs. *NUS1* is yeast homolog of CBP. Arabidopsis CPTs (AtCPT1-9) and tomato CPTs (SlCPT1-7) were numbered according to the published articles (Kera *et al*., 2012; Akhtar *et al*., 2013)

In the phylogenetic analysis, all three *PaCPTs* belong to the cytosolic/ER clade II CPT. At the sub-clade level, with strong statistical supports, *PaCPT1* is clustered with *LsCPT1* (lettuce), *SlCPT3* (tomato), and *AtCPT1* (Arabidopsis), all of which are known to synthesize dehydrodolichyl (Brasher *et al*., 2015; Qu *et al*., 2015; Kwon *et al*., 2016). On the other hand, *PaCPT2* and *PaCPT3* form a close cluster with *LsCPT3* and *TbCPT1-3* (dandelion), known to have implications in NR biosynthesis (Post *et al*., 2012; Qu *et al*., 2015). This phylogenetic tree suggested that guayule possesses two sub-clades of diverged CPTs within the CPT clade II. It can be inferred from these results that *PaCPT1* is likely to be involved in the dolichol metabolism, while *PaCPT2* and *PaCPT3* are involved in NR metabolism.

The phylogenetic tree for CBPs was also constructed. CBP is exclusively found in eukaryotes, and thus the phylogenetic structure of CBP shows a monophyletic group (Figure 1B). The duplications of CBP (i.e., two CBP isoforms) are restricted to lettuce and dandelion in the Cichorieae tribe (within Cichorioideae sub-family under Asteraceae family), and other plants with high quality genome sequences encode a single copy of CBP. Interestingly, PaCBP and other single copy CBPs from the Asteroideae sub-family (sunflower HaCBP and Jerusalem artichoke HtCBP) are tightly clustered with the laticifer-specific CBPs (lettuce LsCBP2 and common dandelion ToCBP2), even though there is no developed laticifer in sunflower and artichoke. Thus, the CBP gene duplication event likely occurred prior to the divergence of the two sub-families and the advent of laticifer in Cichorioideae subfamily.

### Complementation in *rer2*Δ *srt1*Δ yeast

In eukaryotes, dolichol is an indispensable carrier molecule of sugar moiety for post-translational protein modifications. Therefore, all eukaryotes have at least one set of CPT and CBP to synthesize dehydrodolichyl diphosphate (i.e., dolichol precursor), and the absence of either CPT or CBP is lethal. Yeast *(Saccharomyces cerevisiae)* has one homolog of CBP *(NUS1)* and two homologs of CPTs (RER2 and *SRT1).* We previously developed a yeast strain that has double deletions in *RER2* and *SRT1*., in which its lethality is rescued by expressing a plasmid copy of *RER2* (Kwon *et al*., 2016). To examine the ability of PaCPT and/or PaCBP to rescue the dolichol-deficiency of *rer2*Δ *srt1*Δ., these guayule genes were first transformed to this strain individually or in *PaCPT/PaCBP* pairs, and then the URA-selective plasmid containing *RER2* was removed by 5-fluoroorotic acid (5FOA) selection (Figure 2). This enabled us to assess the PaCPT activity in the absence of both of the yeast CPTs.

**Fig. 2.**
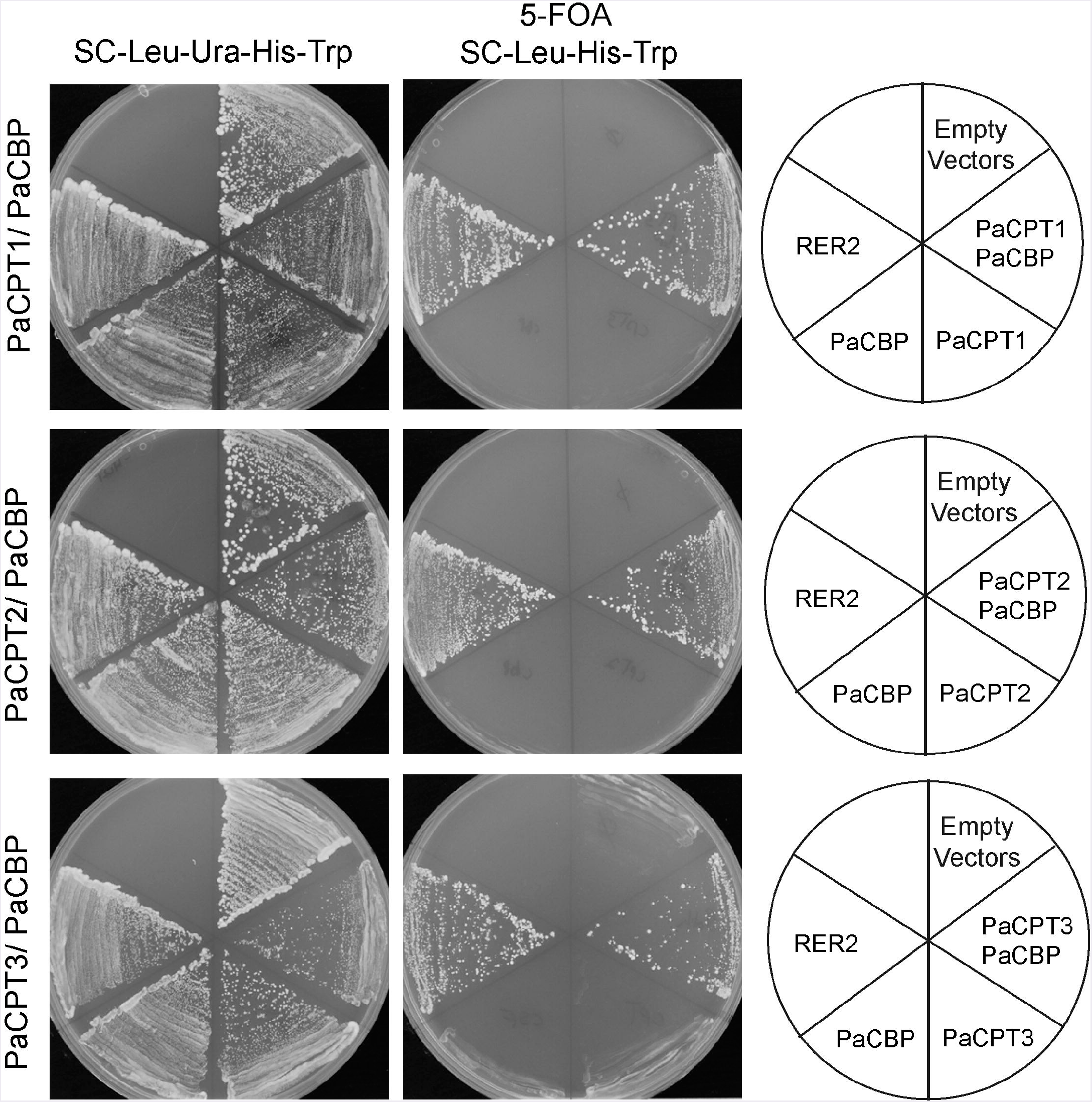
Complementation of *rer2*Δ **srt1**Δ yeast by *PaCPT1-3* and *PaCBP*. The yeast strain, *rer2*Δ *srt1*Δ is lethal but is maintained by expressing *RER2* in URA-selectable plasmid. This strain was used to transform plasmids expressing each *PaCPT1-3* and *PaCBP.* The successful transformants were streaked on 5FOA selection plates to remove *RER2* containing URA-plasmid. Yeast growth in 5FOA selection was observed only by PaCPT/PaCBP pairs or by retransformed RER2 in TRP-plasmid. No growth was observed when PaCPT alone or PaCBP alone was expressed.

It should be noted that if *cis*-polyisoprenes significantly longer than dolichol (C55-C90) were synthesized, or in the absence of CPT enzymatic activity, the *rer2*Δ *srt1*Δ lethal phenotype will not be rescued. In all cases tested, the *rer2*Δ *srt1*Δ strain transformed with empty plasmids or a single plasmid that contains either *PaCBP* or one of the *PaCPTs* failed to grow upon removal of the *RER2-URA* plasmid by 5FOA selection. In contrast, co-expression of *PaCBP* and one of the three *PaCPTs* could effectively complement the lethality of the yeast CPT-knockouts upon removal of the *RER2-URA* plasmid. Our results indicate that each PaCPT can produce isoprenoid molecules analogous to dehydrodolichyl diphosphate only in the presence of PaCBP but not together with endogenous yeast CBP, Nus1.

It is notable that no discernable phenotypic differences were observed in the rescued yeast strains. In principle, if a long carbon NR (~1 million Da) is primarily synthesized by CPT and CBP, the NR should not be able to rescue the dolichol-deficient yeasts. Hence, the robust growth of dolichol-deficient yeast by *PaCPT* and *PaCBP* expression may indirectly suggest that a high molecular weight NR was not synthesized when CPT/CBP were reconstituted */n v/vo* in yeast.

### *In vitro* enzyme assays

The surviving yeast strains after 5FOA selection should be able to bring about necessary CPT activities for the yeast survival in the *rer2Δsrt1*Δ yeast. To assess the guayule CPT activities, microsomes were prepared from the rescued yeast strains from the complementation assay, and radiolabelled ^14^C-isopentenyl diphosphate (^14^C-IPP) and the priming molecule, farnesyl diphosphate (FPP), were co-incubated with the isolated microsomes. When the incorporations of ^14^C-IPP to hydrophobic polymers were measured, significant IPP-incorporation activities ranging from 14.4% to 50.8% incorporation were measured in 2-hour assays (Table 1). Their specific activities were comparable to (in the cases of *PaCPT1/PaCBP1or PaCPT3/PaCBP* activity) or exceeded (in the case of *PaCPT2/PaCBP* activity) the CPT activities previously reported from lettuce CPT and CBP (Qu *et al*., 2015). The CPT activity of the microsomes from the rescued yeast strain transformed with *RER2* as a complementation control was substantially lower (4.5 – 15.6 fold) than those from guayule *PaCPT/PaCBP* co-expression. The lower CPT activity of RER2 control strain is attributable to the imbalance between overexpressed RER2 and its binding partner NUS1, expressed at a native level in yeast.

**Table 1.** Specific activities of PaCPTs and PaCBP in isolated microsomes.

| cDNA expressed | *Activities | *IPP-incorporations |
| --- | --- | --- |
| in *rer2Δ srt1Δ* | (pmol IPP $\mu\text{g}^{-1}$ hr$^{-1}$) | (%) |
| *RER2* | $2.9 \pm 0.4$ | $2.9 \pm 0.1$ |
| *PaCPT1 / PaCBP* | $13.0 \pm 1.7$ | $14.4 \pm 0.5$ |
| *PaCPT2 / PaCBP* | $45.4 \pm 1.3$ | $50.8 \pm 5.0$ |
| *PaCPT3 / PaCBP* | $22.0 \pm 0.4$ | $24.6 \pm 2.6$ |

\*Data are the average ± S.D. from three independent assays.

The resulting ^14^C-labelled products were separated by the reverse-phase thin layer chromatography (RP-TLC) optimized for the separation of dolichol-sized polymers. In the RP-TLC analysis, we detected no obvious differences among the radiolabelled isoprene polymers produced from any of the PaCPT and PaCBP pairs (Figure 3A), and their polymer size was slightly smaller than, but still comparable to, the dehydrodolichols produced by RER2 (Figure 3B). We could not find evidence that the *cis*-isoprene polymer, significantly longer than dolichol, is synthesized by PaCPT and PaCBP in yeast microsomes.

**Fig. 3.**
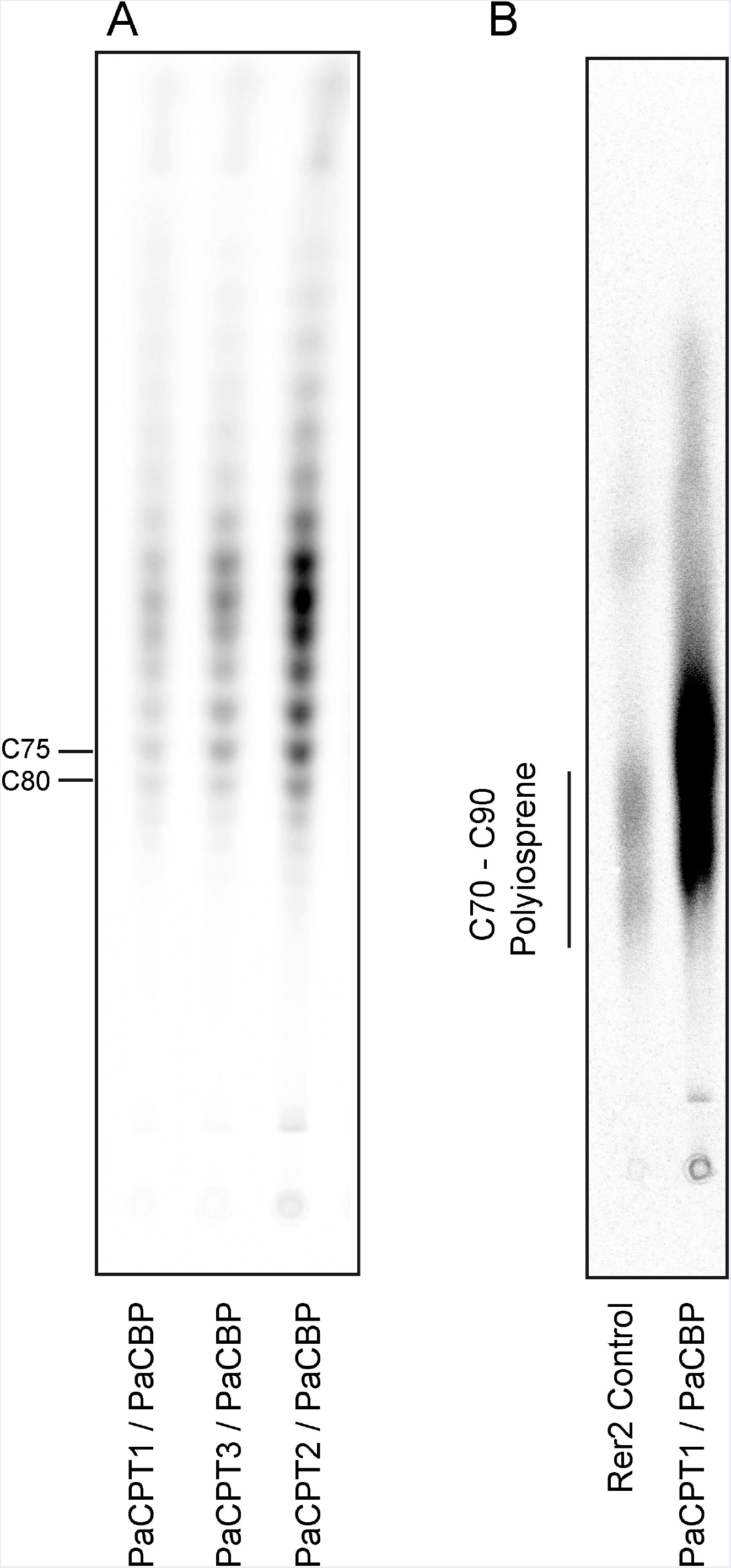
Isoprene product separations by reverse phase thin layer chromatography. Microsomes were prepared from the yeast strains selected on 5FOA in Figure 2, and *in vitro* enzyme assays were performed using the microsomes and ^14^C-isopentenyl diphosphate, followed by RP-TLC of extracted *cis*-polyisoprene products and phosphorimager analysis. A) The **cis*-* polyisoprene product profiles are identical between different PaCPT/PaCBP pairs. B) Dehydrodolichol products from PaCPT/PaCBP are slightly shorter than those from the RER2-containing microsomes. RER2 dehydrodolichol products range in sizes of C70 – C90

### Protein interactions by split-ubiquitin yeast 2-hybrid assay

The complementation of *rer2*Δ *srt1*Δ yeast strain by co-expression of *PaCPT* and *PaCBP*., but not by a single *PaCPT* or *PaCBP* expression, implied that both proteins are required for dolichol metabolism. Thus, possible interactions between each PaCPT and PaCBP were examined by split-ubiquitin yeast 2-hybrid assay developed for protein interactions of membrane-bound proteins (Stagljar *et al*., 1998). In this assay, the restoration of ubiquitin by N-terminal ubiquitin (Nub) and C-terminal ubiquitin (Cub) allows a release of transcriptional regulator to nucleus to activate the auxotrophic marker genes and the β-galactosidase reporter gene. A single residue mutation (I to G) of natural Nub protein (i.e., NubI to NubG) prevents the auto-assembly of the ubiquitin, unless NubG and Cub binding is mediated by the interaction of the proteins of interest each fused to NubG and Cub. Therefore, interactions between PaCPT and PaCBP proteins translationally fused to Cub and NubG can be assessed by yeast growth on selective medium and β-galactosidase activity.

As a control, co-expression of *PaCBP-Cub* fusion and *OST1* (α-subunit of the oligosaccharyltransferase complex localized on the ER)-NubI fusion allows restoration of ubiquitin, thereby presenting efficient yeast growth and strong β-galactosidase activity (Figure 4A). In contrast, changing NubI to NubG in the same experiment did not permit yeast growth, suggesting PaCBP and OST1 do not interact with each other and cannot bring the two ubiquitin subunits together as expected as a negative control (Figure 4A). Similarly, PaCPTs 1-3 were individually fused to NubG, and each fusion construct was co-expressed with PaCBP-Cub to assess whether PaCPT-PaCBP interaction occurs to restore ubiquitin. The interaction between each of the PaCPTs and PaCBP could be clearly inferred by the efficient yeast growth and the by β-galactosidase activities. Interestingly, the interactions were only observed in one orientation of the PaCPT fusion protein (Figure 4B). The NubG-PaCPT orientation allowed activations of the growth markers and reporter gene, whereas the PaCPT-NubG orientation did not show evidence of protein-protein interactions. Wherever interactions were observed, the *HIS3* marker gene was activated very efficiently as yeast could grow at up to 25 mM of AT (3-amino-1,2,4-triazole), a competitive inhibitor of HIS3. In addition, β-galactosidase activities were comparable to that from the positive control. Therefore, the split-ubiquitin Y2H provided molecular evidence that each PaCPT and PaCBP interact with each other to form a hetero-protein complex in yeast.

**Fig. 4.**
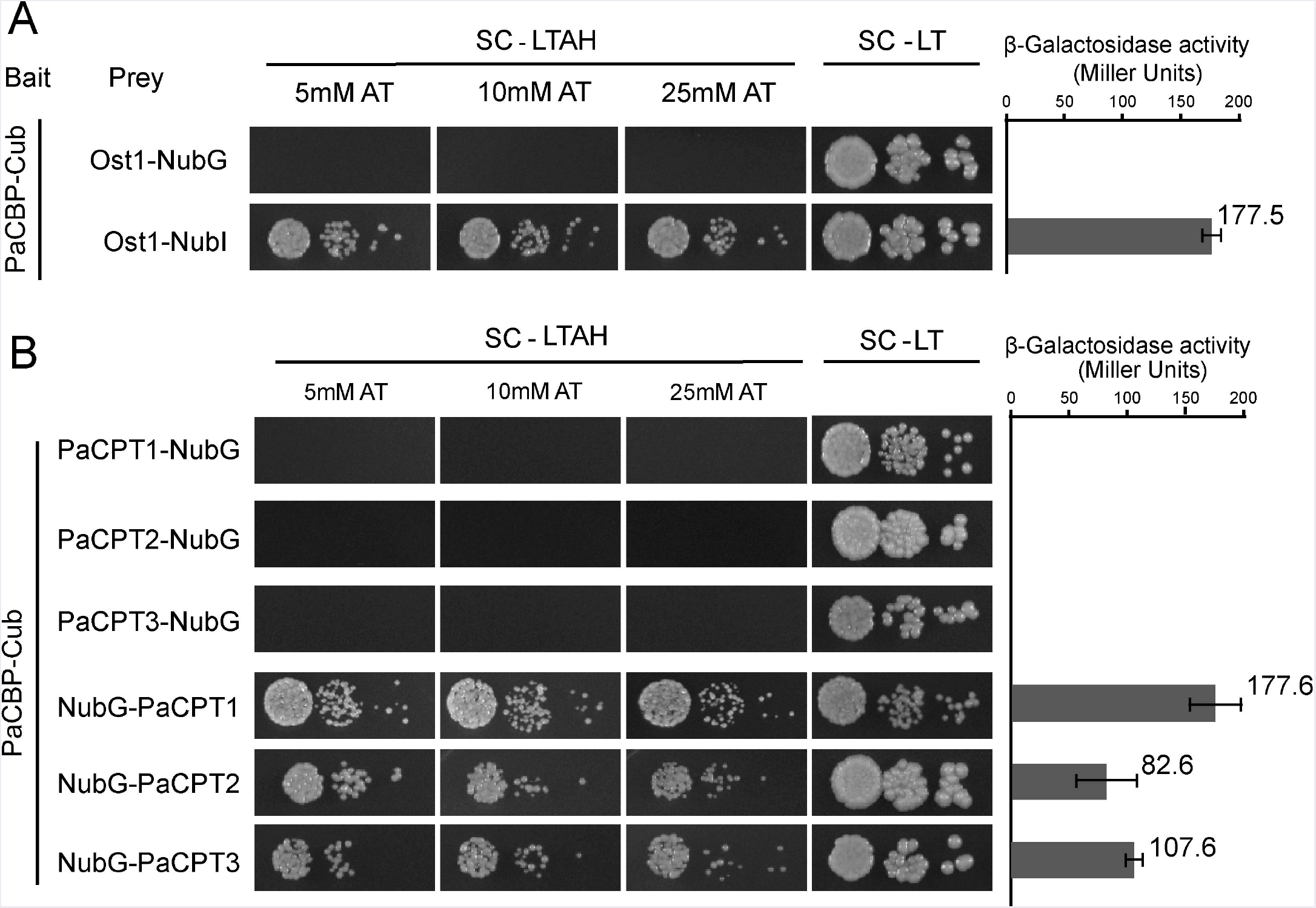
Split ubiquitin yeast 2-hybrid assays to assess the protein interactions between PaCPT1-3 and PaCBP. The bait protein, Cub, was fused at the C-terminus of PaCBP in all assays. Under the selective growth condition, SC-LTAH, yeast growth was observed only when the bait and prey proteins interacted either through NubI-Cub auto-assembly (A: Ost1-NubI, positive control) or through the PaCPT/PaCBP interaction-mediated NubG-Cub assembly (B: NubG-PaCPT3, NubG-PaCPT2, NubG-PaCPT3). PaCPT/PaCBP interactions occurred only when NubG was fused on the N-terminus of PaCPTs. Under the non-selective growth condition, SC-LT, all yeast strains grew regardless of the protein interaction. As an important negative control, NubG-fused Ost1 protein did not interact with Cub-fused PaCBP (A), demonstrating a specific interaction of PaCBP to PaCPT, but not to Ost1. Yeast growth was observed in a wide range of 5 - 25 mM AT, indicating a strong activation of the AT reporter gene by the ubiquitin-activated transcription factor. The β-galactosidase activity was measured in the corresponding yeast strains that showed protein interactions in the absence of AT. PaCPT1/PaCBP interaction showed the same level of β-galactosidase activity as the Ost1-NubI control while PaCPT2/PaCBP and PaCT3/PaCBP showed lower activities.

### Protein interactions by co-immunoprecipitation

In addition to the split-ubiquitin Y2H approach, PaCPT1-3 and PaCBP interaction were independently examined by co-immunoprecipitation assays (Co-IP). PaCPT1-3 and PaCBP were tagged by FLAG- and HA-epitope, respectively, and their proteins were prepared by *in vitro* transcription and translation. Green fluorescent protein (GFP), which has a similar size to PaCBP and the PaCPTs, was used as a negative control in Co-IP experiments. All *in vitro* co-transcribed and co-translated PaCPT/PaCBP proteins were detectable in input by immunoblotting with either FLAG or HA antibody (Figure 5A/B, Input). Incubation of FLAG-antibody immobilized magnetic beads resulted in the co-IP of FLAG-PaCPTs with HA-PaCBP as detected by FLAG-antibody and HA-antibody immunoblotting, respectively, while HA-GFP control protein could not be pulled down by FLAG-PaCPT2. (Figure 5A). Similarly, immunoprecipitation with HA-antibody immobilized magnetic beads resulted in the co-IP of HA-PaCBP with FLAG-PaCPTs while HA-GFP protein could not pull-down FLAG-PaCPT2. (Figure 5B). These results showed that each of the PaCPT1-3 and PaCBP interact with each other to organize hetero-protein complexes when co-expressed *in vitro*.

**Fig. 5.**
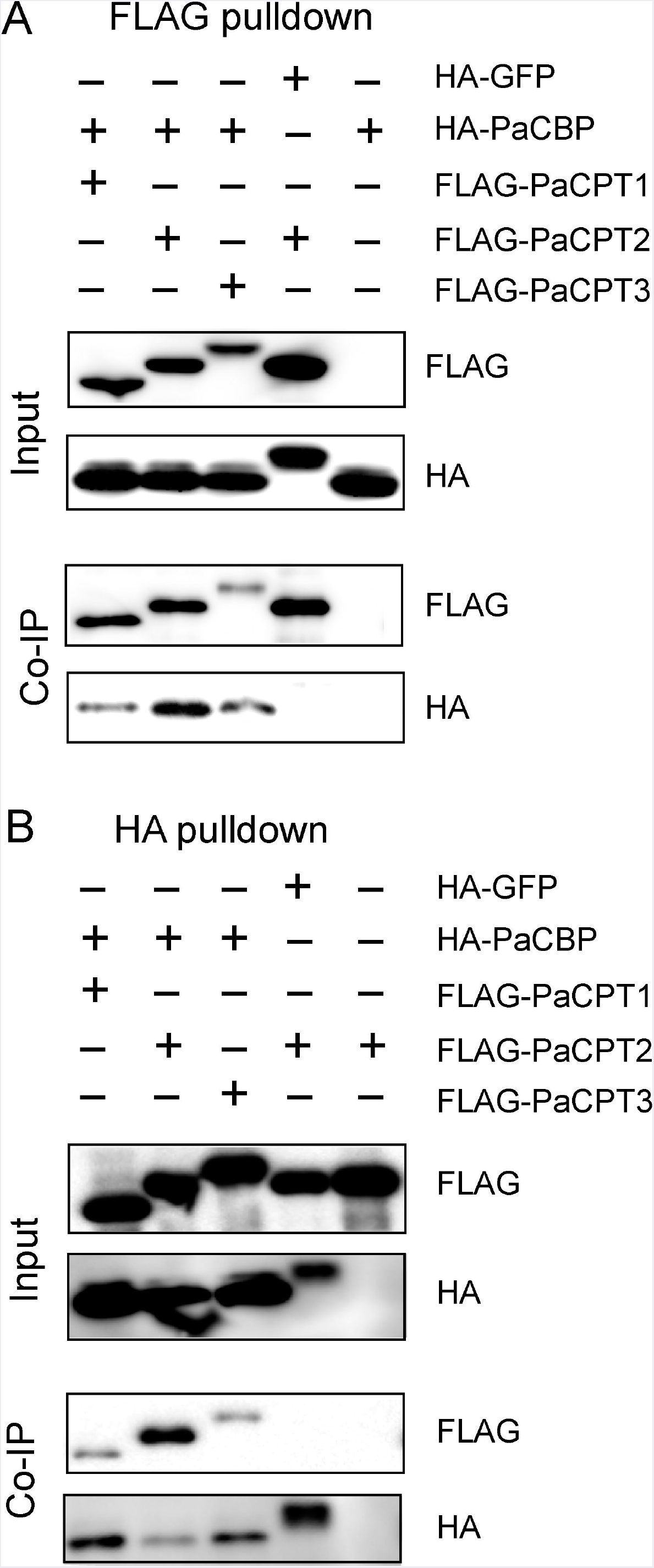
Co-immunoprecipitation of recombinant PaCPT/PaCBP proteins. A) HA-PaCBP could be co-immunoprecipitated by FLAG-PaCPT1-3, and B) FLAG-PaCPT1-3 could be co-immunoprecipitated by HA-PaCBP. PaCPT1-3 and PaCBP proteins tagged with FLAG- and HA-epitopes, respectively, and were prepared by *in vitro* transcription and translation. GFP tagged with an HA-epitope was used as a negative control. The information of the protein mixture is detailed on the top panel with the appropriate epitope-tag. The antibodies used in the immunoblot are indicated beside blots (FLAG: FLAG antibody or HA: HA antibody).

### Expression analysis

Cold temperature is known to induce natural rubber biosynthesis in guayule (Madhavan *et al*., 1989). Based on the positive effect of cold-stress on rubber synthesis, Illumina sequencing of guayule stem and leaf tissues after cold-stress was recently performed, and the sequencing data are publicly accessible at NCBI (short read archive number: SRP107961). Using these data, we conducted a RNA-seq analysis to examine the transcript abundance of *PaCPT1-3* and *PaCBP* (Figure 6).

**Fig. 6.**
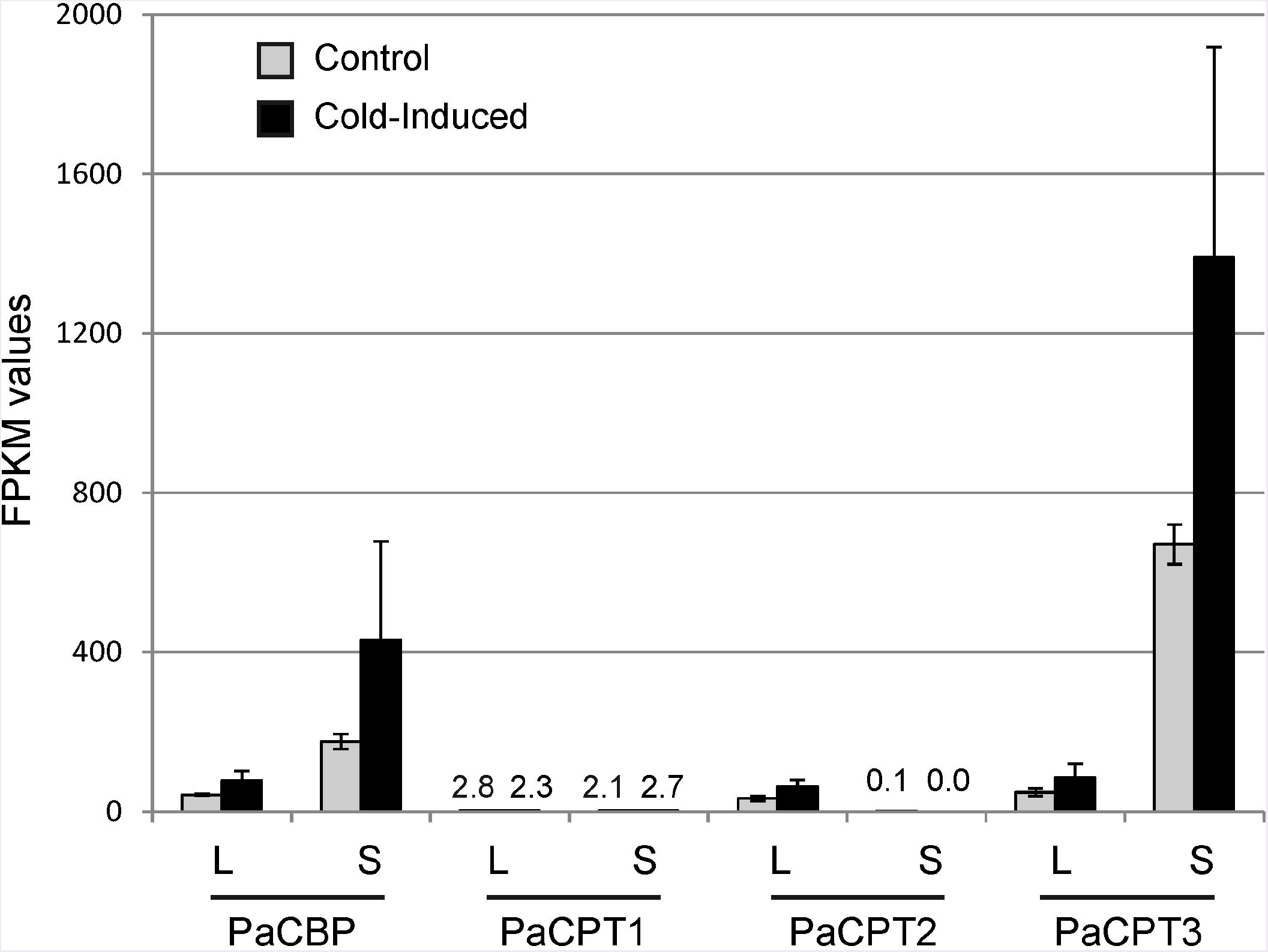
RNA-seq analysis of *PaCBP* and *PaCPT1-3*. FPKM (Fragments Per Kilobase Million) values for *PaCBP* and *PaCPT1-3* were calculated from the publicly available data (short read archive number: SRP107961). L: leaf sample, S: stem sample. Data are means ± S.D. (n=3).

*PaCPT1* transcripts were maintained at a relatively constant basal level in leaves and stems under both normal and cold-treated conditions. These data suggest the basal metabolic role of *PaCPT1* in dolichol metabolism and are congruent to the phylogenetic analysis, in which *PaCPT1* is clustered with other *CPTs* involved in the dolichol metabolism. *PaCPT2* transcripts were hardly detectable in stem tissues while detectable at moderate levels in leaves. Of the three *PaCPTs* examined, *PaCPT3* showed the highest level of expression in stem (2- to 3-orders of magnitude higher levels than *PaCPT1/2*)., and its transcript abundance increased 2.1-fold by cold-treatment in stem. As NR is primarily synthesized in the parenchyma cells of the guayule stem (Kajiura *et al*., 2018), these data suggest that *PaCPT3* is the primary *CPT* that contributes to the NR biosynthesis in guayule stem. On the other hand, transcript levels of *PaCBP* showed a lower degree of fluctuation, compared to *PaCPT1-3* gene family. Its expression levels closely matched those of *PaCPT2* and *PaCPT3* in leaves. Noticeably, the *PaCBP* transcript abundance is highest in stem and is increased by 2.4-fold by cold-treatment, precisely mirroring the expression pattern of *PaCPT3* in stem.

## DISCUSSION

NR is an irreplaceable biomaterial for hundreds of medical and industrial products, but its supply depends entirely on the rubber tree. In nature, more than 7,000 plants are known to produce NR (Rivano *et al*., 2013), but the quality of NR in most plants do not meet the NR polymer properties required for industrial uses. The exceptions are Russian dandelion *(Taraxacum kok-saghyz)*., lettuce *(Lactuca sativa)*., and guayule *(Parthenium argentatum)* as they can synthesize ~1 million average Mw NR. Biochemical and reverse genetic data for NR biosynthesis in lettuce and dandelion have been reported (Post *et al*., 2012; Epping *et al*., 2015; Qu *et al*., 2015); however, little is known about NR biosynthesis in guayule at the level of individual cDNA and enzyme, except for the activity data from the isolated guayule rubber particles (Cornish and Siler, 1995). In this work, we identified and characterized three *PaCPT* isoforms and one *PaCBP.* While the results from this work confirmed some known facts, such as the formation of CPT/CBP protein complex and the lack of NR biosynthesis by CPT/CBP *in vitro*., they also provided additional new insights of NR biosynthesis in guayule.

Although the whole genome sequence has not been completed in guayule, we identified one *PaCBP* from 983,076 reads. Considering the read depth, it is unlikely that any expressed, sufficiently diverged *PaCBP* isoform is present in major tissues of guayule. Thus, *PaCBP* appears to be encoded as a single copy gene in guayule. Similarly, a single copy of *CBP* has been identified from the whole genome sequences and transcriptomics of the rubber tree, *Hevea brasiliensis* (Tang *et al*., 2016). Outside of NR-producing plants, the sequence analyses of high quality plant genomes (e.g., Arabidopsis, tomato, poplar, grapevine, and sunflower) also indicate that a single copy of *CBP* is present in these species. On the contrary, lettuce and dandelion, both originating from the same phylogenetic lineage to the tribe level (Cichorieae tribe), have two isoforms of *CBP* – one being general for dehydrodolichyl diphosphate biosynthesis in all cells and the second *isoform* being specific for NR biosynthesis in the laticifer. This *CBP* gene duplication appears to be restricted to the laticiferous plants from the Cichorieae tribe but is not common in other plants, even including NR-producing plants. Taken together, it is apparent that neo-functionalization and recruitment of a specialized *CBP* isoform are not required for quality NR biosynthesis because regardless of the presence of two isoforms of *CBP*., guayule and rubber tree can all synthesize high quality NR. It can be further inferred that a single *CBP* in the rubber tree and guayule should simultaneously support both primary dolichol and secondary NR metabolism. In such a scenario, we envision a challenge in finding the *in planta* evidence (i.e., RNAi or mutant) showing the necessity of *CBP* for NR biosynthesis in rubber tree and guayule, as severe knock-down by silencing or knock-out of *CBP* will be lethal in these plants.

Among the three *PaCPTs* in guayule, a few lines of evidence support that *PaCPT3* is the key isoform involved in guayule NR biosynthesis. 1) Phylogenetically, *PaCPT2* and *PaCPT3* closely clustered with lettuce *LsCPT3* and dandelion *TbCPTl-3*., whose expressions are specific to laticifer (Post *et al*., 2012; Qu *et al*., 2015), and RNA interference of *TbCPT1-3* resulted in significant NR reduction (Post *et al*., 2012). 2) In our RNA-Seq analysis, the *PaCPT3* transcript level was three orders of magnitude higher than that of *PaCPT1* in stem where NR is known to accumulate. 3) The expression patterns of *PaCPT3* closely coincide with *PaCBP* in two tissues under normal or the induced (cold-stress) condition. Despite clustering closely with *LsCPT3, PaCPT2* had a negligible level of expression in stem. Similarly, *PaCPT1* showed a basal level of expression and clustered phylogenetically with other *CPTs* implicated in dehydrodolichol diphosphate biosynthesis. Collectively, these data indicate that *PaCPT3* supports NR biosynthesis in guayule. To further substantiate this, our future efforts will be focused on silencing or overexpression of *PaCPT3* to observe NR reduction or increase in guayule, respectively.

In addition to identifying the candidate genes necessary for NR biosynthesis in guayule, a suite of molecular and biochemical experiments (i.e., complementation in *rer2*Δ *srt1*Δ yeast, *in vitro* enzyme assays, Y2H, and Co-IP) reliably demonstrated that each of the PaCPT1-3 and PaCBP form hetero-protein complexes to synthesize dehydrodolichyl diphosphate. Despite the formation of protein complexes, we were not able to find biochemical evidence that *cis*-polyisoprenes, longer than dehydrodolichol (dephosphorylated form of dehydrodolichyl diphosphate), can be synthesized using PaCPT3 and PaCBP – the prime candidates for NR biosynthesis in guayule. The polyisoprene products synthesized *in vitro* by recombinant PaCPT3/PaCBP were essentially identical to those from PaCPT1/PaCBP or PaCPT2/PaCBP (Figure 3A). Although speculative, some other proteins or cofactors, energy sources for continued polymerizations, or sophisticated protein nano-structures may be necessary to complete sufficiently lengthy NR biosynthesis*in vitro*.

It was reported that a recombinant rubber tree CPT (HRT1) or a lettuce CPT (LsCPT3) alone without the respective recombinant CBP was sufficient to synthesize NR with average ~1 million Da Mw, when expressed in (or reconstituted with) the washed rubber particles isolated from the rubber tree (Asawatreratanakul *et al*., 2003; Yamashita *et al*., 2016). To date, these two reports, which use washed rubber particles, are the only studies demonstrating *in vitro* reconstitutions of ~1 million kDa NR using recombinant proteins. It is attractive to include the washed rubber particles (protein and lipid mixtures) as structural components *in vitro*., but this complicates the interpretation of the *in vitro* data. In addition, both reports showed no need of externally supplied rubber tree CBP or lettuce CBP for NR biosynthesis despite of their *in planta* interaction (Qu *et al*., 2015; Yamashita *et al*., 2016). Furthermore, these results contradict the *in vivo CBP* silencing evidence in lettuce and dandelion, which independently demonstrated the absolute necessity of *CBP* for NR biosynthesis (Epping *et al*., 2015; Qu *et al*., 2015). The discrepancy of the data cannot be easily resolved unless the rubber tree and lettuce/dandelion have different mechanisms for NR biosynthesis. Further investigation is necessary to address whether NR formation is catalyzed by the same or distinct mechanism in the rubber tree (Euphorbiaceae family) and other NR-producing plants (e.g., Asteraceae family; lettuce, dandelion, guayule, and sunflower).

In summary, three *PaCPTs* and one *PaCBP* cDNAs were identified from guayule in this work, and protein-protein interactions between PaCPTs and PaCBP have been substantiated by *rer2*Δ *srt1*Δ yeast complementation assays, split ubiquitin yeast 2-hybrid assays, and co-immunoprecipitation. The co-expression of each *PaCPT* and *PaCBP* in yeast allowed for the formation of the PaCPT-PaCBP hetero-protein complex for the efficient enzymatic synthesis of dehydrodolichyl diphosphate in microsomal *in vitro* enzyme assays. The RNA-Seq and phylogenetic analyses indicated *PaCPT3* to be the most likely *CPT* involved in guayule NR biosynthesis. Collectively, these data suggest that any future bio-engineering efforts should be directed towards *PaCPT3* and its binding partner, *PaCBP* to increase NR yield in guayule.

## ACKNOWLEDGEMENTS

We thank Professor Igor Stagljar (University of Toronto, Canada) for providing the split ubiquitin membrane yeast 2-hybrid system. This work was supported by the National Science and Engineering Research Council of Canada (NSERC) and Canada Research Chair (CRC) program to D.K.R. This work was also supported by a grant from the Next-Generation BioGreen 21 Program (SSAC, grant number: PJ01326501), RDA, Republic of Korea, and Basic Science Research Program by the National Research Foundation of Korea (NRF) funded by the Ministry of Education (2017R1A6A3A03003409) to M.K.

